# Injectable 3D microcultures enable intracerebral transplantation of mature neurons directly reprogrammed from patient fibroblasts

**DOI:** 10.1101/2023.12.10.570992

**Authors:** Janko Kajtez, Fredrik Nilsson, Kerstin Laurin, Andreas Bruzelius, Efrain Cepeda-Prado, Marcella Birtele, Roger A. Barker, Freja Herborg, Daniella Rylander Ottosson, Petter Storm, Alessandro Fiorenzano, Mette Habekost, Malin Parmar

## Abstract

Direct reprogramming of somatic cells into induced neurons (iNs) has become an attractive strategy for the generation of patient-specific neurons for disease modeling and regenerative neuroscience. To this end, adult human dermal fibroblasts (hDFs) present one of the most relevant cell sources. However, iNs generated from adult hDFs using two-dimensional (2D) cultures poorly survive transplantation into the adult brain in part due to the need for enzymatic or mechanical cellular dissociation before transplantation. Three-dimensional (3D) culturing methodologies have the potential to overcome these issues but have largely been unexplored for the purposes of direct neuronal reprogramming. Here we report a strategy for direct *in vitro* reprogramming of adult hDFs inside suspension 3D microculture arrays into induced DA neurospheroids (iDANoids). We show that iDANoids express neuronal and DA markers and are capable of firing mature action potentials and releasing dopamine. Importantly, they can be gently harvested and transplanted into the brain of a Parkinson’s disease rat model to reproducibly generate functionally integrated neuron-rich grafts. The 3D culturing approach presented here thus eliminates a major bottleneck in direct neuronal reprogramming field and, due to its simplicity and versatility, could readily be adapted as a culturing platform used for a broad range of transplantation studies as well as disease modeling.

## Introduction

Recent breakthroughs in cellular reprogramming approaches have paved the way for direct cell fate conversion of somatic cells such as fibroblasts into induced neurons (iNs) without transitioning through an intermediate proliferative state^1–3^. Moreover, direct reprogramming can successfully be done to a specific neuronal subtype by applying a defined set of transcription factors and fate determinants (e.g., dopaminergic^4,5^, cholinergic^6^, striatal medium spiny^7^, GABAergic^8,9^). As such, this unique approach challenges the traditional views on cell identity and opens new opportunities in biomedical research^10^. For instance, besides being an invaluable tool to create patient and disease specific neurons for modeling of age-associated neurological disorders^11–15^, direct reprogramming could provide a non-pluripotent cell source on-demand for tissue engineering applications and autologous personalized transplantation-based therapies for neurodegenerative diseases. In this respect, adult human dermal fibroblasts (hDFs) represent one of the most relevant biological source material for patient-specific applications of direct neuronal conversion. However, iNs generated from adult hDFs in two-dimensional (2D) cultures poorly survive the transplantation into the adult brain. Instead, most conversion protocols used for transplantation studies have been established using murine fibroblasts^8^ or human embryonic fibroblasts^16^ due to their malleability, high conversion efficiency, and high cell fitness of the generated product. On the other hand, a couple of studies that explored grafting of iNs generated from adult hDFs only when the cells were transplanted into the much more supportive postnatal mouse brain^17,18^. Therefore, successful intracerebral grafting of human iNs directly generated from adult patient skin samples continues to be a major challenge in the field.

One of the inherent shortcomings of 2D cultures is that they require cells to be enzymatically or mechanically dissociated from the substrate before harvesting and transplantation resulting in high cell death. Three-dimensional (3D) culturing methodology has the potential to overcome these issues^19^ and in this respect a recent proof-of-principle study that has shown the generation of iNs inside 3D brain-mimicking hydrogels^20^. However, this study was limited to primary mouse embryonic fibroblasts and bulk hydrogel culture approach that still requires sample dissociation for transplantation.

To address the above-mentioned limitations, we hypothesized that sufficiently small 3D cellular structures could provide the benefits of an *in vivo*-like environment permissive to direct cell-fate conversion while being both readily injectable (without the need for culture dissociation) and protective against mechanical and biological stresses during the transplantation procedure. To test the hypothesis, we developed a strategy for direct *in vitro* reprogramming of adult hDFs inside suspension 3D microculture arrays (Fig. 1a). This approach relies on the ability of fibroblasts to self-assemble into compact spherical structures when devoid of substrate-interactive features. In parallel to cellular self-assembly taking place, lentivirus-mediated expression of a defined set of transcription factors initiates cell-fate conversion of hDFs into dopaminergic (DA) neurons, a potential cell source for modeling and treating of Parkinson’s disease (PD). Using this approach we were able to create thousands of induced DA neurospheroids, here termed iDANoids, without the need for any specialized equipment. We further show that these iDANoids express neuronal and DA markers both at the mRNA and protein levels as well as fire action potentials and release dopamine. In addition, these iDANoids could also be gently harvested without the need for enzymatic or mechanical dissociation and directly injected into the brain of a PD rat model with minimal stress. We found that in contrast to cells reprogrammed in conventional 2D culture, iDANoids survive the transplantation and reproducibly generate healthy neuron-rich grafts. The grafted cells survived long-term with evidence of electrophysiological maturation and functional integration into the host circuitry.

**Fig. 1.**
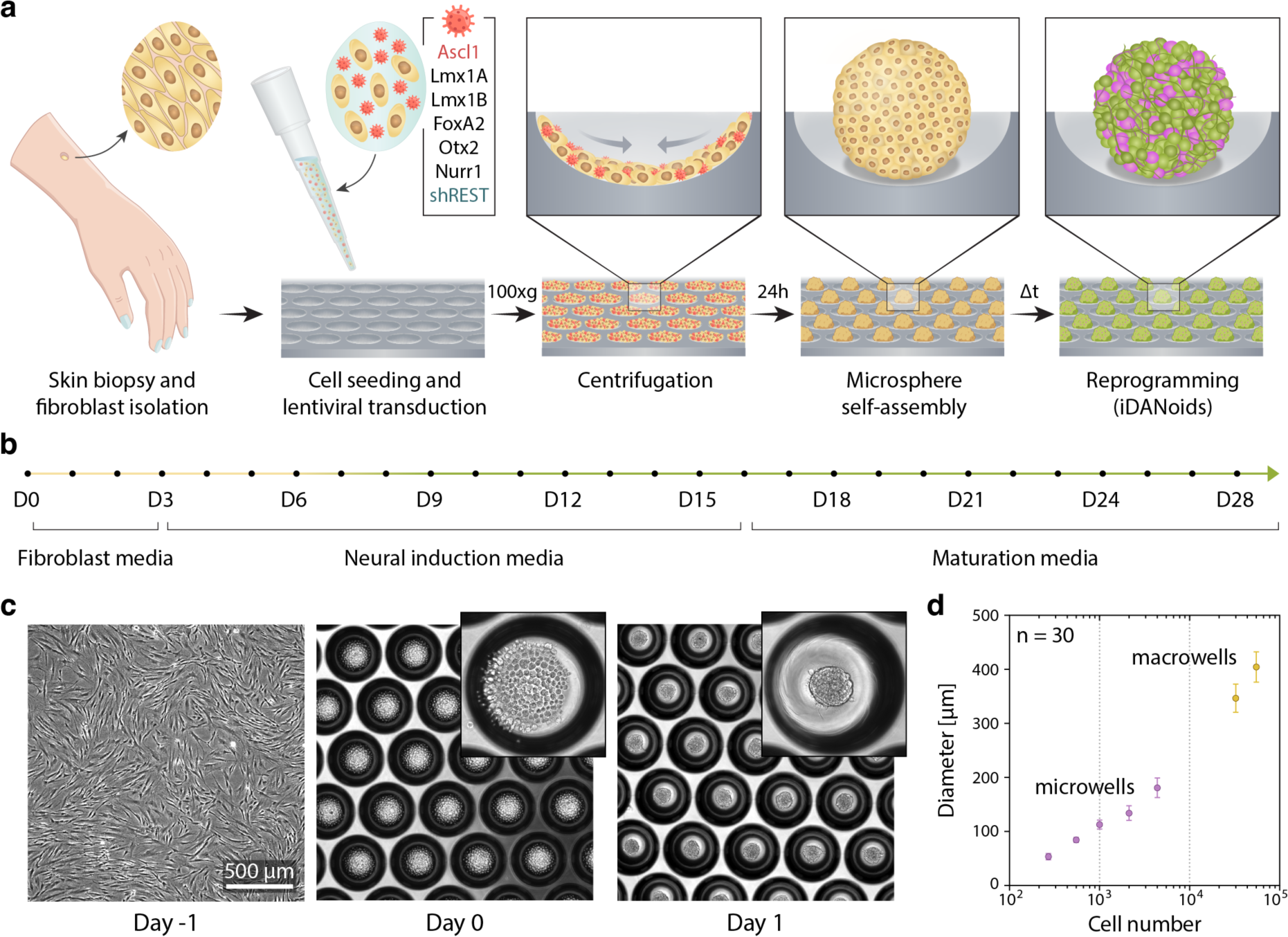
The methodology of iDANoid generation. **a**, Schematic illustration depicting major steps in the developed procedure for direct reprogramming of hDFs in suspension 3D microcultures to generate arrays of iDANoids. **b**, Reprogramming timeline showing media composition used at different stages of the protocol. **c**, Brightfield images showing reproducible self-assembly of hDFs into spheres within 24 h after seeding into arrays of ultra-low attachment conical microwells. **d**, Measurements of sphere diameter 48 h after cell seeding showing our ability to reproducibly tune the size of iDANoids according to the number of cells seeded in each well.

In summary we have created a new 3D culturing approach that eliminates a major bottleneck for direct neuronal reprogramming and due to its simplicity and versatility, could readily be adapted by others as a culturing platform used for transplantation studies as well as for disease modeling, drug screening, and other biomedical applications.

## Results

### Direct neuronal reprogramming of adult hDFs inside suspension 3D microcultures

The suspension 3D reprogramming approach presented here was designed to be compatible with long term *in* vitro culture as well as with standard procedures for intracerebral cell transplantation. As such, it needed to fulfil several requirements: (1) create reproducibly spheroids of a given size; (2) ensure an even distribution of lentivirally delivered conversion factors to ensure homogeneous conversion; (3) maintain spatial separation of individual spheroids during the culturing period without the risk of fusion to form bigger aggregates; (4) enable the gentle harvesting of spheroids for transplantation without the need for cellular dissociation; (5) easy scale-up to allow for high-throughput production able to accommodate larger studies. To achieve this, we seeded hDFs (isolated and expanded from a healthy 61-year-old individual) on an array of conical microwells with ultra-low attachment surface. Lentivirus was mixed with cells during seeding to ensure homogenous exposure. After gentle centrifugation, we observed that cells were distributed evenly between microwells in the array. Within 24 hours, hDFs in each microwell self-assembled into well-defined spherical structures (Fig. 1c). The presence of clear edges to these structures and the lack of wells with individual non-integrated cells indicated that there was a high efficacy to the aggregation process. By varying the seeding cell number, we were able to robustly control sphere diameter (Fig. 1d).

Using this approach, we successfully generated hDF spheres with defined sizes ranging from 68 ± 5 µm to 179 ± 18 µm in diameter corresponding to 250 and 4000 cells per microwell respectively (Fig. 1d). Seeding less than 250 cells in each microwell resulted in inconsistent aggregate formation with compromised structure indicating that a certain number of cells is required for successful and reproducible self-assembly. Conical macrowells were required when seeding more than 4.000 cells for the generation of larger microspheres (e.g., 345 ± 26 µm in diameter for 30.000 cells per sphere). Direct conversion of hDFs into neurons was initiated via expression of six reprogramming factors (Ascl1, Lmx1a, Lmx1b, FoxA2, Otx2, Nurr1) previously identified by us to give rise to a high yield of dopaminergic cells in 2D cultures^14^. Expression of these factors was accompanied by the knockdown of REST complex as shown by us previously for the efficient reprogramming of adult hDFs in monolayers^18^. For the first two weeks, spheres were cultured in neuronal induction medium containing selected small molecules and growth factors known to promote neuronal conversion. Afterwards, maturation media that contained only the growth factors was used until the desired experimental endpoint (Fig.1b).

We performed immunocytochemistry to confirm successful direct neuronal conversion of patient hDF aggregates into iDANoids. The day after seeding, microspheres were positive for the fibroblast marker, vimentin, but not for neuronal markers (Fig. 2a top). By contrast, at day 30 we observed that iDANoids were positive for neuronal markers (MAP2, TAU, TUBB3) and tyrosine hydroxylase (TH, rate-limiting enzyme that catalyzes synthesis of DA precursor L-DOPA) confirming the change in cell identity (Fig. 2a bottom). GABAergic neurons were also observed in the 3D culture (Supplementary Fig. 1a). Importantly, the iDANoid array remained spatially unperturbed with no evidence of fusion to adjacent microspheres (Fig. 2b). Notably, neuronal reprogramming was also successful in large iDANoids with 30.000 cells/sphere (Fig. 2c).

**Fig. 2.**
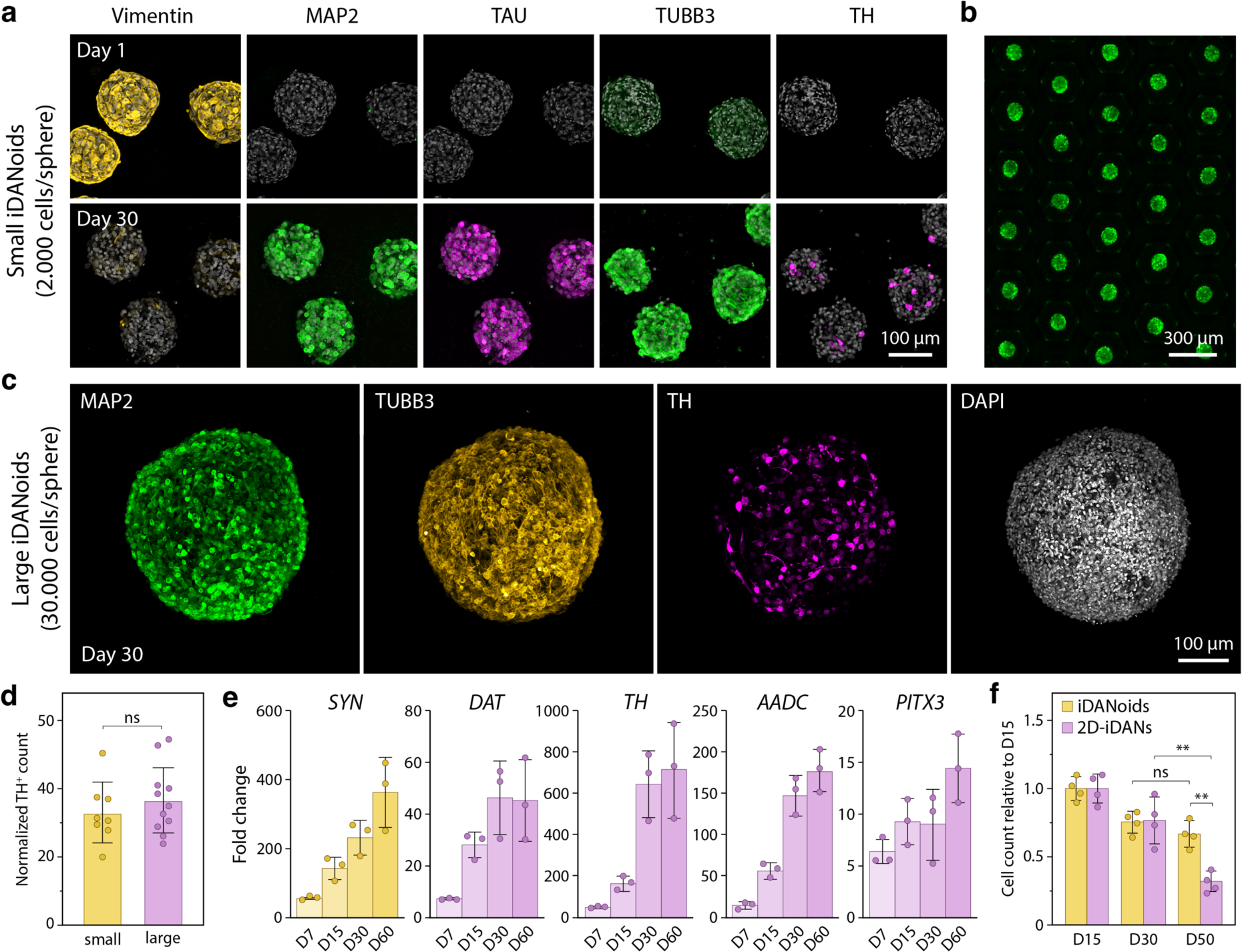
Direct neuronal reprogramming inside suspension 3D microcultures. **a**, Immunocytochemistry of cellular microspheres at day 1 and day 30 after the onset of reprogramming. Maximum intensity projections of fluorescence confocal image stack. **b,** Fluorescence live-cell image showing a segment of a larger iDANoid array at day 30. Cells were labelled with PGK-GFP. **c**, Fluorescence immunocytochemistry of iDANoids seeded with 30.000 cells per microsphere. Maximum intensity projections of fluorescence confocal image stack. **d**, Comparison of DA reprogramming efficacy between small iDANoids (2.000 cells seeded per sphere) and large iDANoids (30.000 cells seeded per sphere) at day 30. The efficacy measurement on the y-axis shows the number of TH^+^ cells normalized to 1000 seeded cells. **e**, RT-qPCR gene expression analysis of iDANoids at different time points. Fold change with respect to the expression levels in the pre-converted fibroblasts. **f**, Cell survival comparison between iDANoids (yellow) and 2D-iDAN cultures at three different time points. Y-axis shows nuclear count for each condition normalized to the mean value at D15.

Next, we compared the conversion efficacy between small (2000 cells/microsphere, ∼120 µm diameter) and large (30.000 cells/microsphere, ∼350 µm diameter) iDANoids to test whether spheroids size affected neuronal conversion. We found no significant differences (Fig. 2d). Progressive maturation of the reprogrammed cells was further investigated by real-time quantitative polymerase chain reaction (RT-qPCR) analysis that demonstrated time-dependent increase in the expression level of mature neuronal markers (*SYN*) and markers associated with DA neurons (*DAT, TH, AADC, PITX3*) (Fig. 2e). Neuronal maturation was also validated at the protein level with NeuN observed at day 50 in iDANoid cultures (Supplementary Fig. 1b). To demonstrate that the reprogrammed neurons can be maintained in the 3D culture for longer periods, we kept iDANoids in culture for 4 months (Supplementary Fig. 1c).

We then compared iDANoids with established 2D reprogramming approaches and observed that there was no significant difference in cell survival at early timepoints, but the robustness of 3D culture became apparent in long term cultures. Thus, at day 50 there were 66 ± 10% surviving cells in 3D cultures while 2D cultures contained only 31 ± 7% surviving cells (Fig. 2f and Supplementary Fig. 1d and 2). These results reinforce a commonly observed issue relating to 2D cultures where cells gradually detach from their plating substrate when maintained long-term while 3D cultures, in contrast, provide increased cell-cell interactions that reduces loss of cells over time. Together, these results indicate that the initial process of cell fate conversion is mainly driven by the expression of the reprogramming factors and is not significantly altered by the change in culturing conditions. However, the enhanced cell-cell contact in 3D culture and lack of dependence on non-physiological substrates, such as tissue culture plastic, plays a major role in the long-term maintenance and survival of reprogrammed cells. Furthermore, it is important to note that the oxygen/nutrient depletion is not a risk factor for the iDANoids used in this study but should be taken into consideration for larger iDANoid cultures (above ∼500 µm)^21^ which might require utilization of a spinning bioreactor^22^.

### Transriptome-wide expression profiling of the iDANoid reprogramming process

We performed bulk RNA-seq and compared the gene expression profiles between hDFs and iDANoids at three different stages of reprogramming (D7, D30, and D50 time points). Principal component analysis (PCA) showed clear transcriptional differences between hDFs, cells in the intermediate reprogramming state (D7), reprogrammed iDANoids (D30 and D50), and iDANs reprogrammed in 2D (Fig. 3a) with most of the variance explained by the reprogramming (first principal component). The difference was least pronounced between D30 and D50 iDANoid samples. Differential expression (DE) analysis of D30 iDANoids with respect to hDFs identified 4333 upregulated and 2867 downregulated genes (Adjusted p-value < 0.01 and |logfc| > 1) (Fig. 3b). Analysis of 14 representative fibroblast-associated genes and 25 neuronal genes confirmed successful conversion of cell identity (Fig. 3c). Notably, analysis of 17 genes enriched in ventral midbrain DA neurons revealed successful DA fate acquisition inside iDANoids starting at the earliest timepoints. In addition to DA neurons, expression of genes related to glutamatergic and GABAergic neurons were also found to increase during reprogramming. Furthermore, analysis of 17 genes associated with neuronal maturation confirmed progressive maturation of the reprogrammed cells.

**Fig. 3.**
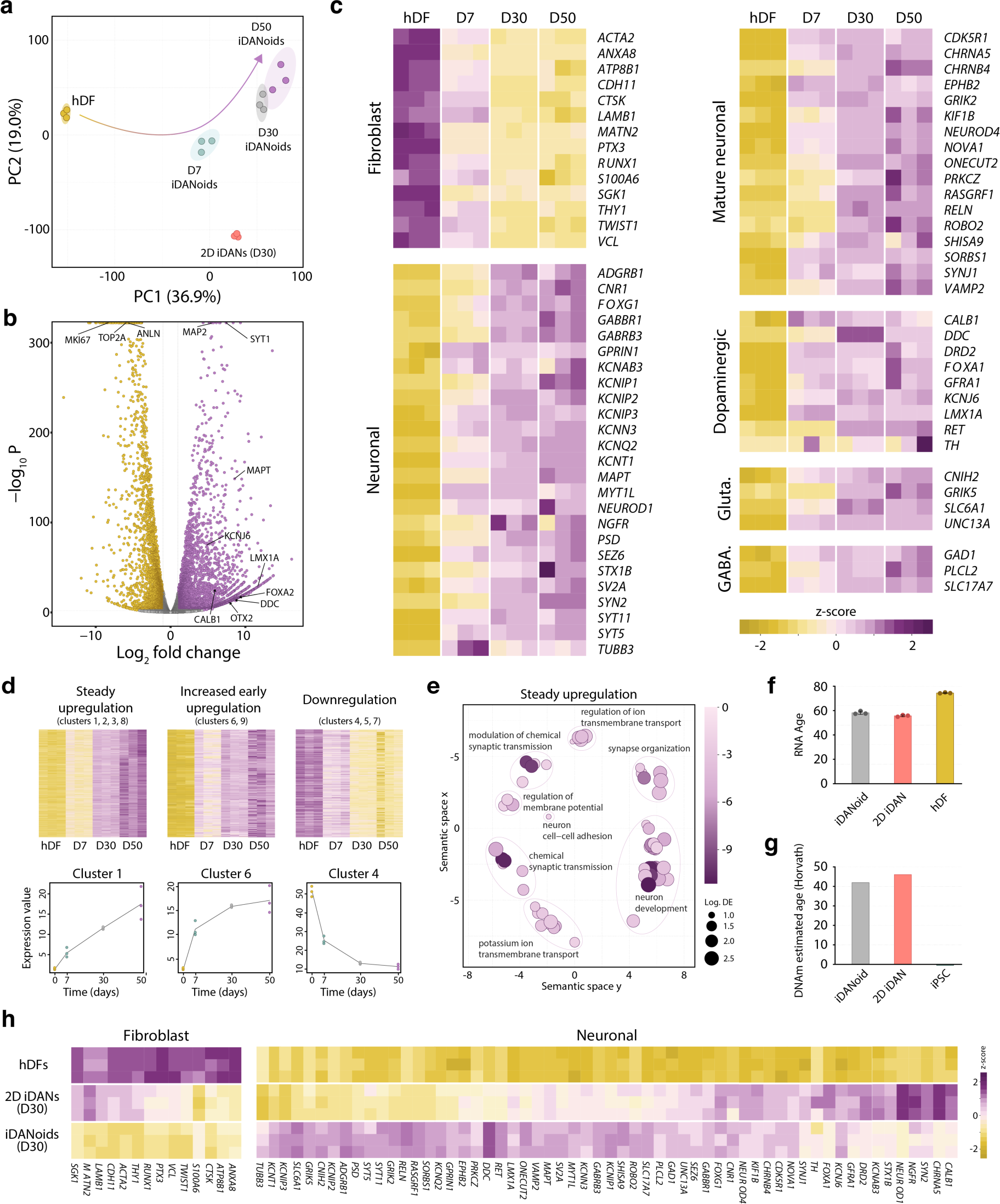
Global gene expression analysis of the reprogramming process. **a**, Principal component analysis (PCA) of transcriptomic profiles obtained from hDFs, iDANoids at 7, 30 and 50 days, and 2D iDANs at day 30 post-transduction. n = 12 data points. Ellipses shown with probability of 0.95. **b,** Volcano plot of differentially expressed (DE) genes (P adj < 0.01) between hDFs and D30 iDANoids. Upregulated DE genes with log_2_ Fold Change (FC) > 1 shown in purple; downregulated DE genes with log_2_ FC < −1 shown in yellow. Selected DE genes labelled with gene symbol. **c,** Heatmap representation of expression levels of selected groups of genes. Values log transformed and scaled by the mean expression of each gene across samples. **d**, Heatmap representation of three groups of genes segregated according to their temporal expression profile clustering using maSigPro analysis. Trajectory plots for representative clusters from each group are displayed below heatmaps. **e**, Gene ontology (GO) analysis of biological processes overrepresented in an upregulated gene set. The scatterplot shows the representative clusters of GO terms obtained from GOseq analysis, while the terms are scattered based on their semantic similarities on the x- and y-axis as determined by REVIGO analysis. Bubble color indicates log_10_ P value while bubble size indicates log_10_ of the number of DE genes in the GO category. **f**, Estimated transcriptional age of iDAnoids and 2D iDANs at D30 in relation to hDFs. **g**, Epigenetic age as predicted by DNA methylation profiling of iDANoids and 2D iDANs at D30 in comparison to iPSCs derived from the same fibroblast line. **h,** Heatmap representation of expression levels of selected groups of genes for hDFs and reprogrammed cells in iDANoids and 2D iDANs at D30.

Next, we applied a two-step regression strategy (maSigPro) to identify clusters of significant genes with similar expression patterns along the experimental time-course^23^. The analysis yielded 9 clusters with different temporal expression dynamics (Supplementary Fig. 3). These clusters were then grouped together according to distinct features identified in the trajectory plots. This resulted in three groups of DE genes. The first group (1804 genes) contained clusters characterized by steadily increasing upregulation over the experimental time course. These clusters could be identified by small changes in the expression value rate of change (slope of the line in the trajectory plot) between the three timepoints (Fig. 3d left). The second group (792 genes) was characterized by an “early on” pattern and steeper increase within the first week of reprogramming compared to other two timepoints (Fig. 3d middle). The third group (1108 genes) contained clusters that were downregulated over the experimental time course (Fig. 3d right). Gene ontology (GO) analysis revealed that the genes upregulated steadily over the experimental time course are related to neuronal terms such as synapse organization, synaptic transmission, and regulation of membrane potential (Fig. 3e) while the genes with a steeper upregulation during the first week related to fate change (Supplementary Fig. 4). GO analysis of the downregulated genes revealed terms such as collagen metabolic processes, actin filament-based processes, response to wounding, and blood vessel development confirming the loss of fibroblast-related properties (Supplementary Fig. 4). Notably, GO analysis revealed there was an upregulation of genes related to neuron cell-cell adhesion, an important aspect of 3D culture, while genes related to cell-substrate interaction were downregulated. When compared to a conventional 2D conversion protocol at D30, iDANoids displayed a more complete switch to a neuronal cell identity. For example, while 14 fibroblast-associated genes were robustly downregulated in iDANoids, 2D iDANs show this comparable level of downregulation only in ∼30% of these genes (Fig. 3h left). Similarly, we observed more consistent upregulation across a panel of 58 genes associated with neuronal identity in iDANoids compared to 2D conversion (Fig. 3h right). Since it is now also well established that aging markers are retained during the process of direct fibroblast-to-neuron reprogramming in 2D, we investigated if the same holds true during direct neuronal reprogramming in 3D. To this end, analysis of bulk RNA-seq data using a transcriptional age calculator correlated reasonably well with the donor’s chronological age and did not reveal a major shift in transcriptional age profile between 2D iDANs (55.6 ± 0.7) and iDANoids (58.1 ± 1.3) (Fig. 3f). No major changes in estimated age between 2D iDANs and 3D iDANoids was observed at the epigenetic level either (prediction based on the amount of DNA methylation using Horvath’s clock) while epigenetic rejuvenation was confirmed in induced pluripotent stem cells (iPSCs) generated from the same fibroblast cell line (Fig. 3g).

### Functional *in vitro* assessment of iDANoids

To validate that the generated neurons were functionally active *in vitro*, we performed whole-cell patch clamp electrophysiological recordings in intact iDANoids at 60 days post induction. A proportion of recorded cells (2/7) displayed the ability to induce action potential upon depolarization (Fig. 4a) along with fast-inactivated inward and outward currents upon membrane depolarization, typical of voltage gated sodium channels and delayed potassium currents (Fig. 4b). In these cells membrane capacitance (Fig. 4c) and AP properties (Fig. 4d) indicated a maturing neuronal function. Taken together, we show that converted cells within iDANoids develop synaptic compartments necessary for function and are able to fire APs upon membrane depolarization.

**Fig. 4.**
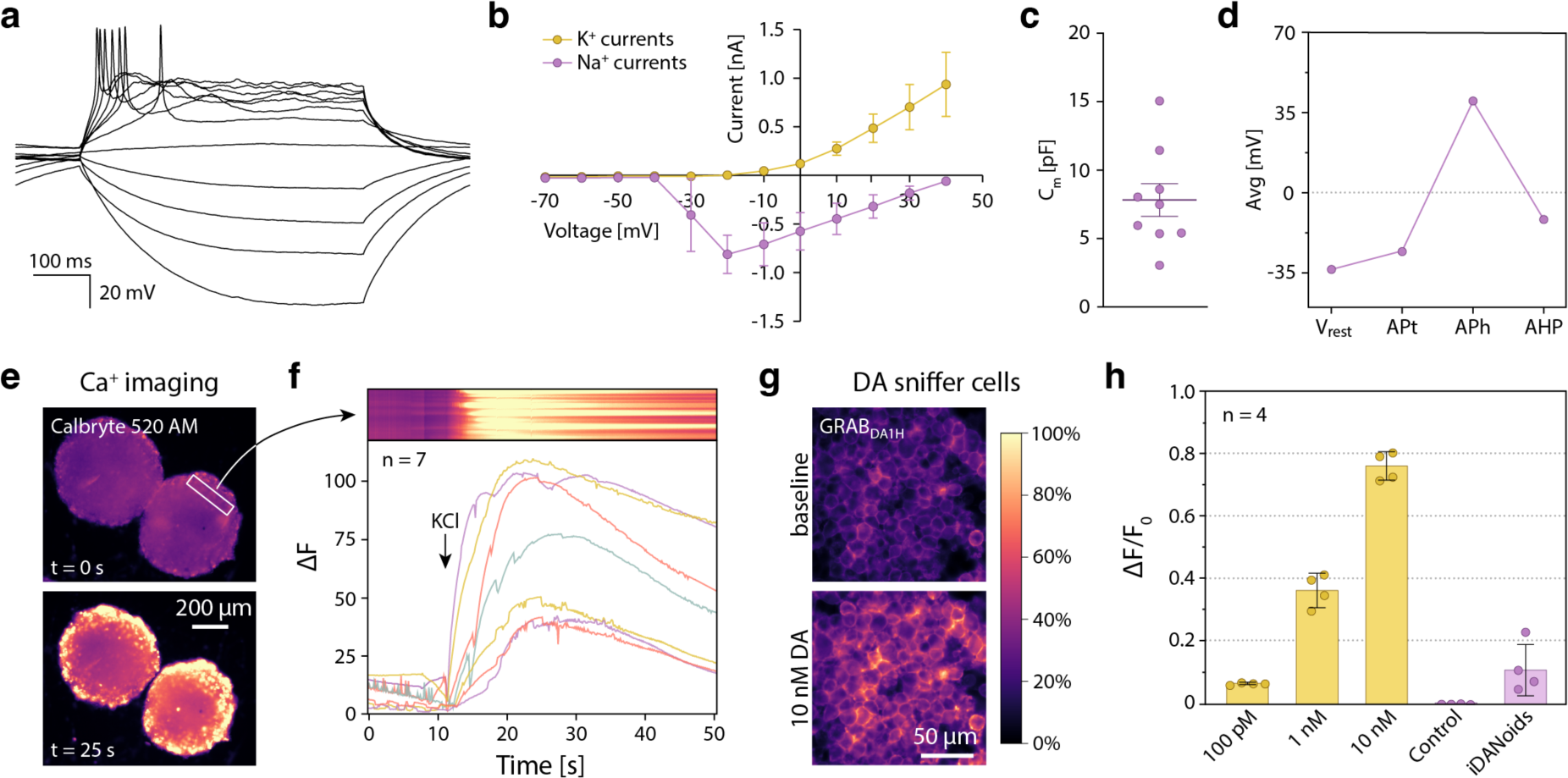
Functional investigation *in vitro*. **a**, Representative trace of evoked action potential from a recorded iDANoid cell. **b**, Inward Na^+^ and outward K^+^ currents plotted against stepwise voltage induction at day 60 (n = 2 cells). Values are presented as mean ± SEM. **c**, Measured capacitance of recorded iDANoid cells. **d**, AP properties, resting membrane potential (V_rest_), AP threshold (AP_t_), AP amplitude (AP_h_) and after-hyperpolarization (APH). Each dot representing the mean value (n = 2). **e**, Fluorescence images of intracellular calcium before and after stimulation of iDANoids with KCl. **f**, Kymograph (top) showing fluorescence intensity change within the selected ROI and traces of change in fluorescent intensity over time for whole iDANoids. **g**, Representative images of GRAB_DA1H_ sniffer cells before and after stimulation with 10 nM DA solution. **h**, Quantification of fluorescent intensity for increasing doses of DA solution and KCl stimulated iDANoids as well as non-reprogrammed hDF spheres (control).

We used calcium imaging to show that reprogrammed cells respond to potassium chloride (KCl) stimulation, a process that causes membrane depolarization and activation of voltage-gated channels in neurons^24^. iDANoids exhibited sharp increase in calcium influx upon KCl addition thus confirming the ability of the reprogrammed cells to respond to chemical stimulus (Fig. 4e,f). To investigate whether iDANoids are capable of releasing dopamine, we took advantage of a microscopy-based detection approach that utilizes dopamine sniffer cells expressing genetically encoded GRAB_DA1H_ DA fluorescent sensor^25,26^. We first determined the sensitivity and the detection range of the sniffer cells *in vitro* by exposing them to increasing concentration of dopamine solution. Sniffer cells showed dose-dependent increase in fluorescence for dopamine concentrations between 100 pM (ΔF/F_0_ = 0.05 ± 0.01) and 10 nM (ΔF/F_0_ = 0.76 ± 0.05) (Fig. 4g). We then triggered dopamine release from iDANoids through KCl stimulation and the solution from depolarized neurons was then harvested, added on top of sniffer cells, and their response was recorded using fluorescence microscopy. Dopamine from iDANoids could be detected (ΔF/F_0_ = 0.07 ± 0.01) at a level corresponding to ∼100 pM (Fig. 4h).

### Generation of induced assembloids

Direct neuronal reprogramming inside suspension 3D microcultures, in contrast to traditional 2D cultures, provides the opportunity for fusion and functional integration of different cell types. To demonstrate this, we placed an iDANoid in contact with a non-dopaminergic induced neurospheroid directly reprogrammed from hDFs via overexpression of two neural conversion genes *Ascl1* and *Brn2* together with REST knockdown (Fig. 5a,b)^18^. To determine the optimal timepoint for sphere fusion, we placed spheres in contact with one another at different days after the initiation of neural reprogramming. D7 spheres already showed signs of interaction after a few hours in contact (5% fused, n = 60). After 3 days, 83% of these pairs fused and after 5 days there was a 98% success rate (Fig. 5c). D14 spheres showed less plasticity with only 75% (n = 35) of the pairs of spheres fusing successfully after 5 days in contact. D21 spheres showed insufficient ability to interact with each other with no fused pairs after 5 days of contact (n = 8). Based on this data, we therefore proceeded with fusing the spheres 7 days after the initiation of reprogramming. Two weeks after fusion, induced assembloids displayed a rich neuronal network with neurons crossing the interface between the two fused spheres (Fig. 5d left). Almost all DA neurons were still located on the iDANoid side of the induced assembloid indicating that reprogrammed neurons do not migrate significantly from their starting sphere to the adjacent sphere, at least over this time frame (Fig. 5d right). This an important since the assembloid platform should not become a homogenous mix but rather a regionally segregated entity that would allow investigation or manipulation of the defined regions. At the same time, the assembloids need to display functional connections between the regions. To show that individual spheres can form functional connections upon fusion, we took advantage of monosynaptic tracing technology based on retrograde transfer of glycoprotein deleted (ΔG) rabies virus across functional synapses ^27,28^. At D0, one set of spheres (termed *source* in the assembloid) were transduced with lentivirus containing the rabies helper construct under the synapsin promoter. At D7, these spheres were fused with spheres that did not contain the rabies helper construct (termed *target*). Then, at D21 ΔG rabies virus was added to the culture. This results in expression of nuclear GFP and cytoplasmic mCherry in *source* neurons. At the same time, retrograde transmission of rabies virus across active synapses would induce expression of cytoplasmic mCherry (but not GFP) in any *target* neurons that form functional presynaptic contacts directly with *source* neurons (Fig. 5e). Immunocytochemistry performed on traced D28 assembloids confirmed functional connections between the fused spheroids (Fig. 5f). While *source* neurons were positive both for nuclear GFP and mCherry, we identified mCherry^+^/GFP^−^ cells with intricate neuronal morphology throughout the target side of the induced assembloid (Fig. 5g).

**Fig. 5.**
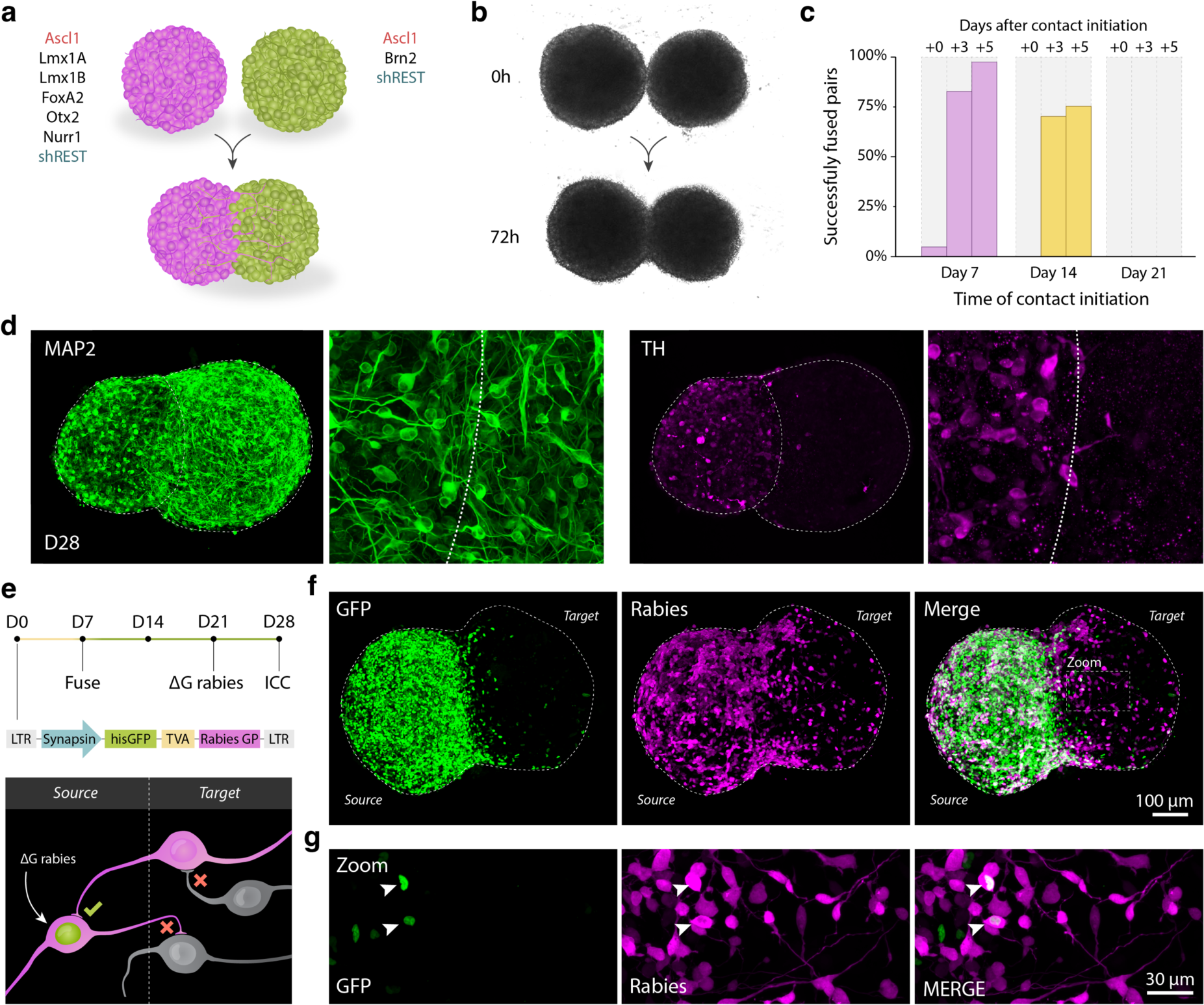
Induced assembloids. **a**, Schematic illustration of the concept behind the generation of induced assembloids. Individual microspheres first undergo direct neuronal reprogramming and then they are fused together. **b**, Brightfield images showing fusion of two D7 microspheres within 72 hours after they are placed in contact. **c**, Bar graph displaying success rate of microsphere fusion with respect to the day of first contact (D7, D14, D21) and time after contact initiation (+0, +3, +5). **d**, Confocal fluorescence image of neuronal marker MAP2 and dopaminergic marker TH in induced assembloid at day 28 after the initiation of conversion. Spheres were fused at day 7. The image displays a maximum intensity projection from a 219 µm thick optical slice. **b**, Schematic illustration displaying the experimental setup and underlying principles of transsynaptic retrograde rabies tracing experiment. **f**,**g**, Representative confocal fluorescence images displaying functional synaptic connections between two spheres in an induced assembloid. The source neurons (GFP^+^/mCherry^+^) are present in the left half of the assembloid that was transduced with the rabies helper construct. Non-transduced right assembloid half shows traced neurons (GFP^−^/mCherry^+^) that formed synaptic connections with the neurons in the adjacent source sphere.

### Intracerebral transplantation into adult rat brain

One of the main goals of our work was to enable the successful engraftment of neurons directly converted from adult hDFs into the adult brain environment. To test whether iDANoids are suitable for this, we took advantage of a well-established preclinical 6-hydroxydopamine (6-OHDA)-lesioned adult PD rat model commonly used to assess dopamine cell replacement therapy approaches. We followed the standard procedure for cell transplantation, by placing the iDANoids into the dopamine denervated dorsolateral striatum and analyzed the outcome 4 weeks post-transplantation (Fig. 6a). iDANoids with 2000 cells/sphere were gently harvested from their microwells and taken up into glass capillaries that are routinely used for intracerebral injection of stem cell-based cell suspensions (Fig. 6b)^29^. The use of a narrow capillary (∼150 µm inner diameter) is important as it minimizes the invasiveness of the procedure and minimizes the trauma of the transplant procedure itself.

**Fig. 6.**
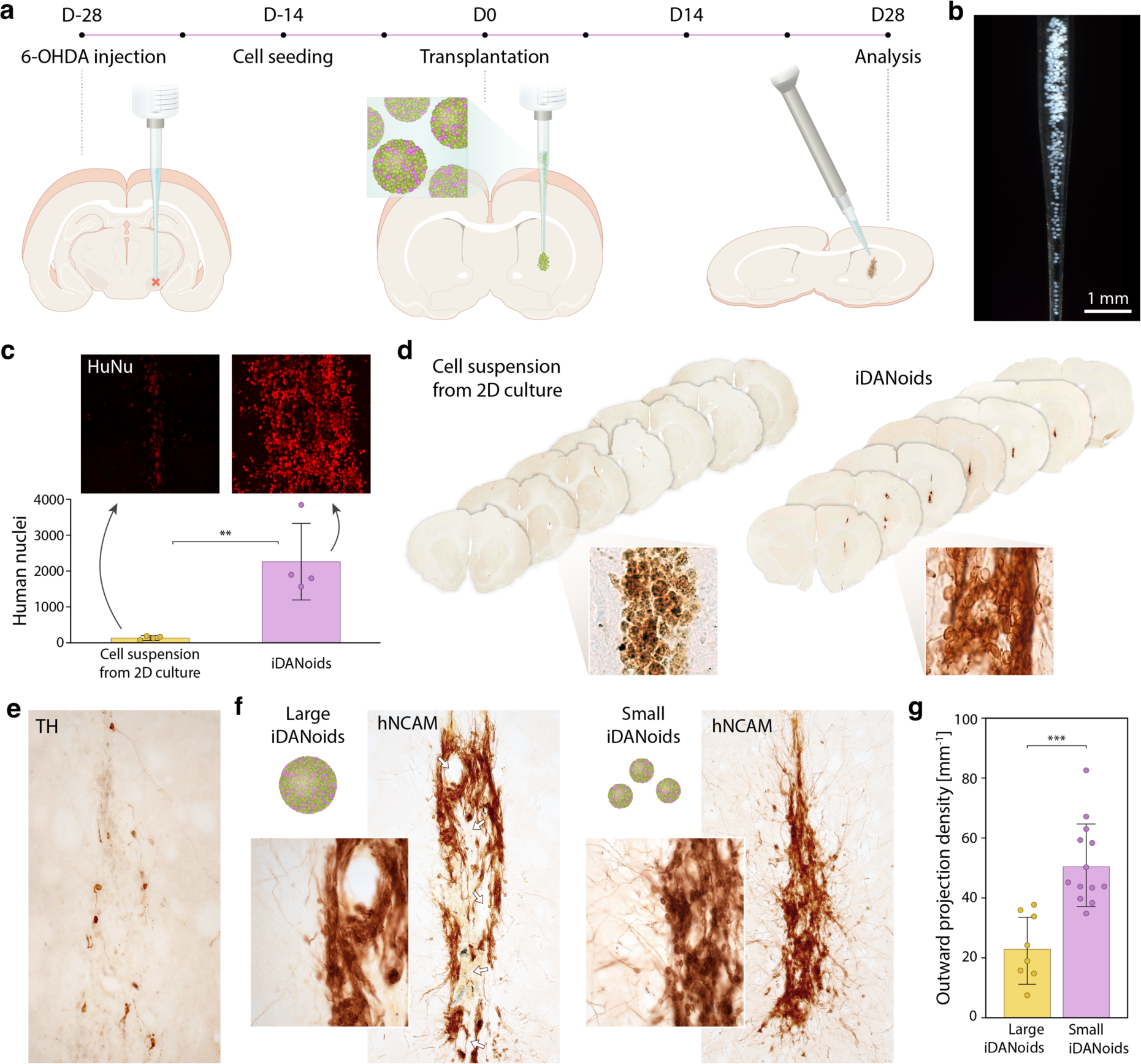
Intracerebral transplantation of iDANoids into adult rat brain. **a**, Graphical representation of the experimental timeline. **b**, Image showing iDANoids collected in a glass capillary, prepared for minimally invasive transplantation into the adult rat striatum. **c**, Count of surviving human nuclei in brain slices 4 weeks after transplantation comparing iDANoids with transplanted cell suspension from reprogrammed cells in adherent 2D culture. Representative fluorescence images accompany the graphs. Counting performed in 1 slice out of 8 from the series. **d**, Images of a series of consecutive rat brain slices with DAB staining for human -specific neuronal marker hNCAM showing graft size and cellular morphology for the two conditions. **e**, DAB staining for TH showing surviving induced DA neurons. **f**, Comparison between grafts from large (30.000 cells/sphere) and small (2000 cells/sphere) iDANoids 4 weeks post-transplantation. For both cases, the same number of cells was transplanted. **e**, Analysis of outward projection density shows that grafting of small iDANoids yields a significantly larger number of projections that grow out into the host tissue.

In the first experiment, we compared the survival of hDFs converted inside traditional 2D cultures and hDFs converted inside microwell arrays to form iDANoids. Fluorescent immunostaining for human nuclei (HuNu) in transplanted brain slices revealed a significant increase in survival of human cells kept in suspension 3D microculture (2278 ± 1066 counted nuclei) in comparison to enzymatically harvested cells from 2D culture where graft survival was minimal (107 ± 69 counted nuclei) (Fig. 6c). Immunostainings for the human neuronal marker hNCAM indicated even bigger differences between the two conditions. While 2D culture yielded a small number of unhealthy-looking cells lacking neuronal morphology, transplanted iDANoids generated neuron rich grafts that span multiple brain sections with compact cell bodies and thin projections emanating into the host tissue (Fig. 6d). DA neurons (TH^+^) were also found inside the graft indicating survival of this neuronal population considered particularly fragile (Fig. 6e).

In the next experiment, we compared the transplantation outcome of small (2000 cells/sphere) and large (30.000 cells/sphere) iDANoids. Large iDANoids require the use of a bigger capillary that leads to more host damage at the time of grafting, and they also produce suboptimal grafts. Grafts from larger iDANoids were riddled with pockets devoid of cells which most likely arise from a low packing capability that leaves empty spaces between adjacent spheres. Furthermore, visibly fewer neuronal fiber outgrowths extended from these grafts into the host tissue (Fig. 6f). In fact, measurements of outward projection density confirmed that smaller iDANoids yield grafts with 2.5 more projections per millimeter of graft circumference than larger iDANoids (Fig. 6g).

### Host integration and long-term survival

To assess the functional properties of the transplanted hDF-derived neurons we performed whole-cell patch-clamp recordings on coronal acute brain slices at 60 days post-transplantation. hDF-derived neurons were labelled with GFP for identification (Fig. 7a). In total, 15 GFP-positive cells were patched for intrinsic membrane properties and demonstrated resting membrane potential (V_rest_) of −31.7 ± 7.90 mV, an input resistance (R_i_) of 4.1 ± 2.3 MOhm, and a capacitance (C_m_) of 7.38 ± 2.49 pF which indicated different stages in functional maturity that was much improved from the *in vitro* condition (Fig. 7b,c). This was further confirmed in their intrinsic firing properties with 40% of the cells were able to generate multiple action potentials (APs) indicative of a neuronal maturity (Fig 7d,e), another 40% only fired single AP indicative of immature state, and 20% of the cells showed no AP in response to steps of depolarization (Supplementary Fig. 5a). Further analysis of the action potential properties revealed an AP threshold of −23.1 ± 8.56 mV, AP amplitude of 40.70 ± 5.03 mV, and afterhyperpolarization of 12.93 ± 2.12 mV (Fig 7b). These properties resemble those of cultured human adult fibroblast-derived neurons ^9,18^, including low maximum sodium (80.1 ± 14.2 pA) and potassium (441.9 ± 76.6 pA) currents evoked by increasing steps of depolarization in voltage clamp mode (Fig. 7f). Tetrodotoxin (TTX) blockade further confirmed the sodium channels activation (Supplementary Fig. 5b). Interestingly, we observed spontaneous APs in 20% of recorded cells (Figure 7g), typical of dopaminergic neurons, that has not yet been observed *in vitro* ^30^.

**Fig. 7.**
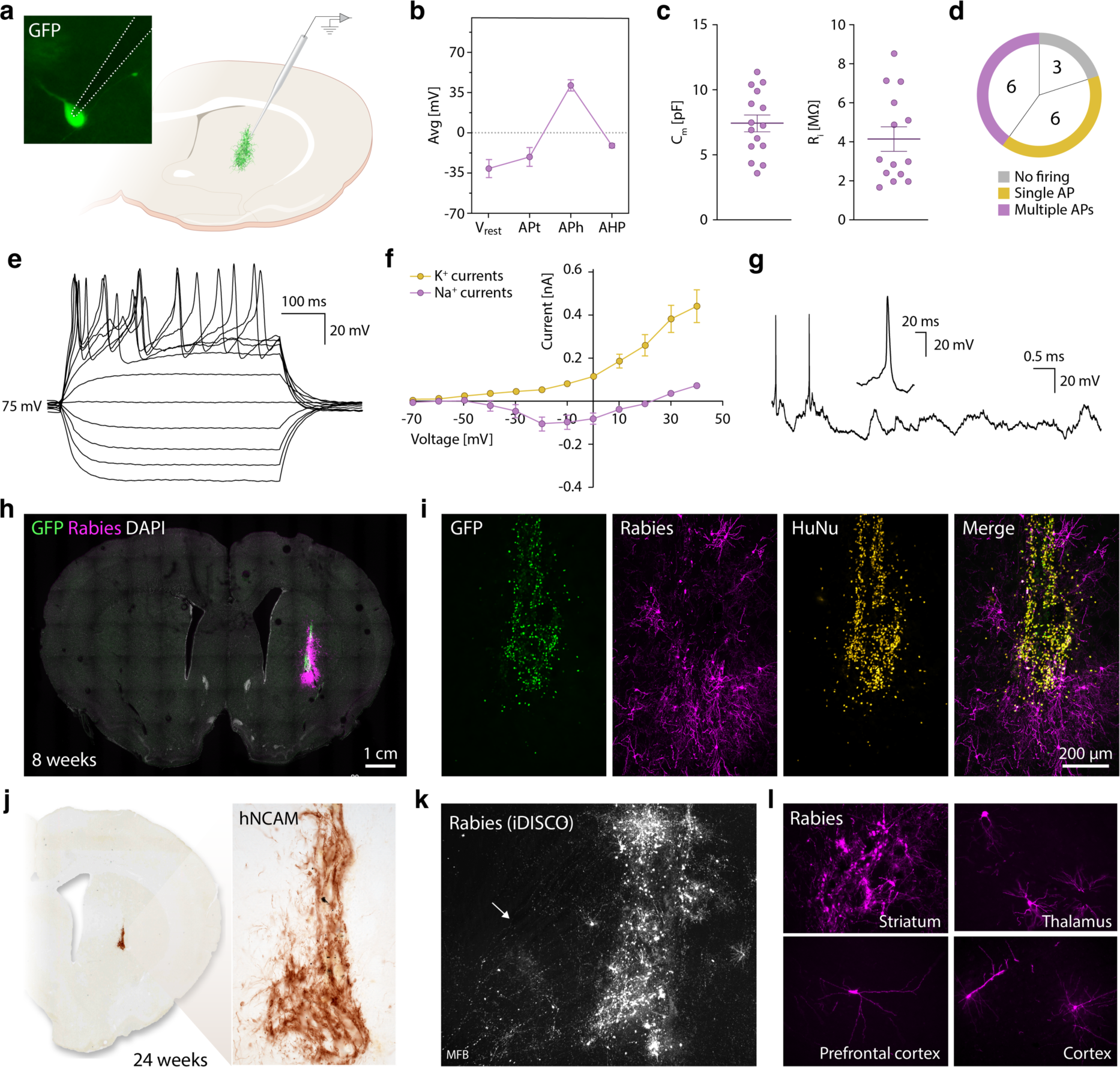
Post-transplantation assessment of functional activity and host integration. **a**, Graphical illustration and fluorescent image showing the micropipette tip accessing a GFP-labelled neuron in an acute brain slice of rat dorsolateral striatum. **b**, Properties of the first observed action potential evoked by the rheobase current injection step such as resting membrane potential (V_rest_), AP threshold (APt), AP amplitude (APh), and afterhyperpolarization. **c**, Summary of capacitance (C_m_) and input resistances (R_i_) of transplanted iDANoid neurons. **d**, Proportion of recorded cells firing none, single, and multiple APs **e**, **f**, Representative voltage responses to depolarizing and hyperpolarizing currents. **g**, Example of spontaneous action potentials firing recorded in a current clamp at holding potential of −70 mV. **h**, Fluorescent confocal image of a whole brain slice showing an iDANoid graft, 8 weeks post grafting. Transplanted cells are marked with nuclear GFP. Host cells making synaptic connections with the graft are labelled with (ΔG) rabies virus (magenta). **i**, Close-up fluorescent image of the same graft with the addition of HuNu staining confirming that the GFP-labelled cells are of human origin. **j**, Overview image and a closeup of hNCAM-DAB staining in a striatal brain slice. **k**, Maximum intensity projection from a series of fluorescent light sheet images showing rabies-traced host-to-graft synaptic inputs. Anatomical landmarks were identified from endogenous background signal. **l,** Fluorescent confocal images of rabies-traced neurons in different brain regions 24 weeks post-transplantation.

To assess functional integration of transplanted iDANoids into the host circuitry, we took advantage of the same monosynaptic rabies tracing technology used to analyze induced assembloids. First, at the time of seeding into microwells, hDFs were transfected with lentivirus containing rabies helper construct. Then, a week before the animals were sacrificed for analysis, they were injected with ΔG-rabies virus at the location of the graft. This results in expression of nuclear GFP and cytoplasmic mCherry in grafted cells. Retrograde transmission of rabies virus across active synapses would induce expression of mCherry (but not GFP) in any host neurons that form presynaptic contacts directly with grafted human induced neurons. Fluorescence staining of brain slices 8 weeks post-transplantation confirmed functional host-to-graft connectivity (Fig. 7h). Nuclei positive for GFP were also positive for HuNu, confirming that these are human neurons originating from grafted iDANoids. A network of mCherry^+^/GFP^−^ neurons with complex morphology surrounded the graft indicating that neurons directly reprogrammed from adult hDFs were not only able to survive the transplantation but also received functional synaptic inputs from the host (Fig. 7i).

Finally, we assessed the potential of human induced neurons from transplanted iDANoids to survive over long periods of time (at least 24 weeks) while maintaining communication with host circuitry. hNCAM-DAB stainings of striatal brain slices from rats sacrificed 24 weeks post-transplantation show neuron rich grafts with dense neuronal projections, sign of morphological maturation of the transplanted cells (Fig. 7j). We then performed iDISCO tissue clearing and light sheet microscopy of whole rat brains with grafts labelled for monosynaptic rabies tracing (Fig. 7k). The results not only show synaptic inputs from the local host environment but also functional connections from distant neuronal populations (e.g., neurons extending projections through the MFB). Closer analysis of immunostained brain slices revealed endogenous neurons from several brain regions including thalamus, prefrontal cortex, and cortex to form long-distance connections with the grafted cells in addition to striatal host circuitry (Fig. 7k).

## Discussion

Increasingly sophisticated stem cell-based 3D culturing techniques for neuroscience applications have shown great potential to overcome many of the inherent limitations of the traditional 2D approaches and to mimic aspects of the native tissue such as spatial organization, cell-cell interactions, mechanical properties, and extracellular environment^31,32^. However, novel culturing platforms for direct neuronal reprogramming remain largely unexplored^20^. Here, we present a robust 3D culturing approach for high-throughput generation of iNs from adult hDFs via direct cell fate conversion inside suspension microculture arrays. Using this approach, we were able to generate large numbers of directly reprogrammed neurospheroids with controlled size and preserved structural integrity.

Major efforts in the field to improve direct neuronal conversion in 2D cultures have been directed towards developing ever more complex combinations of transcription factors, miRNAs, and small molecules for more efficient generation of type- and subtype-specific neural cells. While we focused on the generation of DA neurons inside iDANoid microcultures, the presented approach is highly versatile and could easily be adapted to conversion strategies already established in 2D for direct reprogramming of fibroblasts to other neuronal (e.g., cholinergic^6^, striatal medium spiny^7^, GABAergic^8,9^) and non-neuronal cell types (e.g., astrocytes^33–35^, oligodendrocytes^36–38^).

Our approach offers several advantages over 2D conversion protocols. It does not rely on artificial cell-substrate interactions which are well known to be unstable over long culture periods (leading to cell detachment and loss), but rather on reinforced cell-cell interactions that facilitate long-term survival and cell maintenance *in vitro*. Notably, the combination of tissue clearing and confocal imaging presented here could be translated towards high throughput applications for automated screening experiments^39^. The free-floating nature of the microculture allows for the easy manipulation of individual microspheroids at any given time during the reprogramming process. This allowed us to create induced assembloids, 3D co-culture platform that could pave the way for novel *in vitro* studies of interregional brain communication, neuron-glia interaction, pathology transfer, and disease modeling in a similar manner to that which has been done using stem cell-based brain assembloids^40,41^. Our experiments further reinforced findings that cellular organization into self-supported 3D structures mitigates mechanical and biological stresses and by so doing increases cell survival, preserves cell function, and facilitates host integration after transplantation^42–44^. Most importantly, we used the new approach with great success to transplant iDANoids reprogrammed from adult hDFs into adult rat model of PD. The transplanted cells survived, integrated, and functionally matured in the brain to a level that has not been demonstrated before and presents an important achievement towards *in vivo* studies using patient-specific iNs and potential cell replacement therapies based on direct cell reprogramming.

In summary we have developed a novel suspension 3D microculture approach that has a widespread potential for disease modeling and transplantation using direct neuronal reprogramming of patient-specific fibroblasts.

## Materials & Methods

### Cell lines

hDFs used in this study were obtained from the Parkinson’s Disease Research Clinic at the John van Geest Centre for Brain Repair (Cambridge, UK) and used under full local ethical approvals: REC 09/H0311/88 (University of Cambridge). The subjects’ consent was obtained according to the declaration of Helsinki. For biopsy sampling information see ^18^.

### Cell culture

hDFs were cultured on uncoated T175 flasks in fibroblast medium (DMEM, 10% fetal bovine serum, 1% penicillin-streptomycin) at 37 °C and 5% CO_2_. All media was changed every 2-3 days. Upon reaching a 90% confluency the cells were split with 0.05% trypsin in DPBS and further expanded, frozen down or replated for reprogramming experiments. Fibroblasts re-plated for reprogramming in 2D were seeded onto poly-ornithine, laminin, and fibronectin-coated plates at a density of 25.000 cells per cm^2^ according to our previous study^45^. The following day, the media was replaced with fresh fibroblast media containing the virus at an MOI of 5 per construct. To ensure a more homogenous transduction of cells in 3D cultures, the cells were seeded together with the virus in ultra-low attachment plates: for 250-5000 cells/sphere in Elplasia plates (Corning) were used, for more than 5000 cells/sphere Costar U-bottom ULA 96-well plates (Corning) were used. Two days later, the media was changed to early neuronal conversion medium consisting of NDiff227 supplemented with CHIR99021 (2μM), SB431542 (10μM), Noggin (0.5μg/mL), LDN1931189 (0.5μM), VPA (1mM), LM22A4 (2μM), GDNF (2ng/mL), NT3 (10ng/mL) and db-cAMP (0.5mM). Partial media changes were carried out 2-3 times per week until day 16 when the media was changed to late neuronal conversion media made up of NDiff227 supplemented with LM22A4 (2μM), GDNF (2ng/mL), NT3 (10ng/mL) and db-cAMP (0.5mM). Media was partially changed every 2-3 days until the end of the experiment.

### Lentivirus production

The lentiviral particles were produced in HEK-293 cells according to previously published protocol^46^. In brief, HEK-293T cells were transfected with three helper packaging plasmids along with the plasmid of interest and polyethylenimine (PEI) to facilitate transfection. After 48 hours the media was filtered, ultracentrifuged and resuspended in DPBS. The titer was determined by RT-qPCR using primers against a reference gene (albumin) and a virus specific gene (WPRE) and comparing the expression to a reference virus with an already measured titer.

### In vitro whole cell patch clamp

Electrophysiological recordings were performed 60 days post induction on intact spheres transferred into the recording chamber at experimental timepoint. The chamber was filled with Krebs solution gassed with 95% O2 and 5% CO2 at RT. The composition of the Krebs solution was (in mM): 119 NaCl, 2.5 KCl, 1.3 MgSO4, 2.5 CaCl2, 25 glucose and 26 NaHCO3 with the pH adjusted to 7.4. Spheres were recorded in a free-floating state using a Multiclamp 700B amplifier (Molecular Devices, San Jose, CA, USA) with borosilicate glass pipettes (3–7 MOhm) backfilled with the following intracellular solution (in mM): 122.5 potassium gluconate, 12.5 KCl, 0.2 EGTA, 10 Hepes, 2 MgATP, 0.3 Na3GTP, and 8 NaCl adjusted to pH 7.3 with KOH. pClamp 10.2 (Molecular Devices, San Jose, CA, USA) was used for data acquisition with the current filtered at 0.1 kHz and digitized at 2kHz. Directly after successful break in, the resting membrane potential was measured in current-clamp mode, followed by current injection to stabilize cells at - 60 to −70 mV. To elicit evoked action potentials, current was injected from −20 pA to +35 pA with 5 pA increments. Measurements of inward sodium and delayed rectifying potassium currents was done with cells clamped at −70 mV in voltage clamp mode using voltage-depolarizing steps supplied for 100 ms at 10 mV increments. Data was processed and analyzed with the Clampfit 10.3 (Molecular Devices, San Jose, CA, USA) and Igor Pro 8.04 (Wavemetrics, Portland, OA, USA) software combined with the NeuroMatic package^47^.

### Calcium imaging

iDANoids were incubated for 30 min at 37 °C with 3 µM Calbryte 520 AM calcium indicator (AAT Bioquest) in maturation medium containing 0.02% Pluronic F-127 (Sigma Aldrich). Cells were then rinsed with baseline buffer containing 1.2 mM MgCl_2_, 2 mM CaCl_2_, 150 mM NaCl, 5 mM KCl, 5 mM glucose, and 10 mM HEPES buffer and imaging was performed on an inverted fluorescence microscope equipped with a 20x objective. Stimulated calcium influx was recorded with an exposure time of 100 ms by inducing cell membrane depolarization through the addition of stimulation buffer containing of 1.2 mM MgCl_2_, 2 mM CaCl_2_, 5 mM NaCl, 150 mM KCl, 5 mM glucose, and 10 mM HEPES buffer. Images were analyzed in ImageJ (NIH) and plotted in Prism (GraphPad).

### DA sniffer cells

The HEK-293 Flp-In T-Rex cell line ^26^, stably expressing fluorescent G protein-coupled receptor-based DA sensor GRAB_DA1H_ ^25^, was cultured in DMEM supplemented with 10% FBS, 15 µg/mL blasticidin, and 200 µg/mL Hygromycin B. Sniffer cells were passaged at least twice before they were used in experiments. 48 h before the measurements were taken, sniffer cells were seeded in imaging chambers (8-well, ibidi) coated with poly-L-ornithine and the expression of GRAB_DA1H_ sensor was induced with 1 µg/mL tetracycline. Live DA sniffer cell imaging was performed on a widefield Leica microscope using a 20x NA 1.4 objective. First three images were obtained to determine baseline fluorescence. Then, samples collected from depolarized iDANoids (or non-reprogrammed spheres as a control) were added to the imaging well and three more images were taken to determine the fluorescence response of the sensor. Images were averaged and analyzed in ImageJ (NIH) and results plotted in Prism (GraphPad).

### Immunocytochemistry

The cells were washed twice with DPBS and fixed with 4% PFA (10 min and 30min for 2D and 3D cultures, respectively) in RT and washed twice before adding blocking solution (5% serum, 0.1% Triton) for 1 h in order to prevent non-specific binding. Next, the cells were left in blocking solution with primary antibodies in 4 °C overnight. The following day the cells were washed twice with DPBS, incubated in blocking solution for 30 min in RT followed by secondary antibody (1:200 Jackson ImmunoResearch Laboratories) in blocking solution along with DAPI (1:500) for 1 hour for 2D cultures and overnight for 3D cultures. Lastly the cells were washed twice and left in 4 °C until analysis.

### Clearing of 3D cultures

Once the 3D cultures had been stained, they were cleared in order to improve the imaging quality and enable imaging of the entire sphere. The spheres were put in 20, 40, 60, 80 and 100% methanol for 10 min each. Next, they were left in a DCM-methanol mixture (2:1) for 1 hour and then in 100% DCM for two, 10 min periods. Lastly, the spheres were cleared with ethyl cinnamate and transferred to 96 Well Black µ-Plates (Ibidi, #89626).

### RT-qPCR

The cells were washed twice with DPBS, lysed and total RNA collected using the RNeasy Micro Kit (QIAGEN#74004) according to the manufacturer instructions. RNA (1μg) was reverse transcribed to cDNA with the Maxima First Strand cDNA Synthesis Kit (Thermo Fisher #K1642). The cDNA was mixed with the primers of interest and with SYBR Green Master mix (Roche #04887352001) using the Bravo instrument (Agilent) and analyzed by a LightCycler 480 II instrument (Roche) utilizing a 2-step protocol with a 95 °C, 30 second denaturation step followed by a 60 °C, 60 second annealing/elongation step for a total of 40 cycles. Samples were analyzed with the ΔΔCt-method, normalized against the two housekeeping genes ACTB (actin-beta) and GAPDH (glyceraldehyde-3-phosphate dehydrogenase) and the data presented as a fold change of expression compared to non-converted hDFs. Table S1 contains the list of primers used in the study.

### Bulk RNA sequencing

RNA was extracted using the RNeasy mini kit (Qiagen) following the manufacturer’s instructions. cDNA libraries were prepared using the Illumina TruSeq library preparation kit and sequenced with 2 x 150 bp paired end reads on an Illumina NextSeq 2000 using the P3 flow cell (∼37M reads per sample). Raw base calls were demultiplexed and converted into sample specific fastq files using default parameters of the bcl2fastq program (v 2.20, Illumina). Trimmed reads (TrimGalore, 0.6.1) were aligned to the human genome version GRCh38 (Ensembl release 99) using STAR^48^ (v2.7.3) and quantified using Salmon^49^ (v0.14.0).

A principal component analysis (PCA) was performed through singular value decomposition of a data matrix that had been both centered and scaled using *prcomp* function in R. Identification of differentially expressed genes from gene-level count tables was performed using DEseq2^50^ (2.11.40.7+galaxy1). Heatmaps were generated to visualize gene expression levels by log transforming and mean scaling transcript per million (TPM) values using the *heatmap.2* function in R. Assessment of significant gene expression profile differences was achieved using maSigPro^51^ (1.49.3.1+galaxy1) on gene-level count tables. Functional analysis of differentially expressed genes, selected based on a false discovery rate-adjusted P value (P adj, using Benjami-Hochberg procedure) cutoff of 0.05, was conducted using Goseq^52^ (1.44.0+galaxy0). The outcomes of this analysis were visually summarized utilizing the REVIGO^53^ tool.

RNA age for each sample was estimated using the *predict_age* function in the R package RNAAgeCalc^54^.

### DNA methylation profiling

DNA was extracted from ∼1000 iDANoids (containing 2000 seeded fibroblasts each) or from a T25 flasks (1.2 × 10^6^ fibroblasts plated) using DNeasy Blood and Tissue Kit (Qiagen) following the manufacturer’s instructions. Bisulfite conversion of DNA samples was prepared using the EZ DNA Methylation kit from Zymo Research. Methylation analysis of bisulfite pretreated DNA samples was done using the Illumina Infinium MethylationEPIC v2.0 BeadChip (EPIC). Illumina Genome Studio software was used to perform sample quality control. We calculated individual epigenetic ages for each sample using Horvath’s clock using the R package *cgageR* (version 0.1.0, https://github.com/metamaden/cgageR) after preprocessing using minfi^55^. This method relies on the methylation data from 353 CpGs to produce a continuous value that mirrors an individual’s epigenetic age.

### Animals

All experiments involving live animals were carried out in accordance with the European Union Directive 2010/63/EU and were approved by the Swedish Department of Agriculture and the Lund University ethics committee. Experiments lasting 8 weeks or less were carried out in adult female Sprague-Dawley (SD) rats from Charles River Laboratories. To keep the SDs immunosuppressed and avoid graft rejection they were given daily intraperitoneal injections of Ciclosporin A (10mg/kg) two days prior to cell grafting until the end of the experiment. For experiments lasting more than 8 weeks, adult female athymic rats from Envigo were used instead. All rats weighed at least 225g before the start of any experiment and their weights were recorded every week to keep track of potential weight-loss. All rats were housed in ventilated cages under 12-hour light/dark cycles with free access to food and water.

### Surgeries

All surgeries were carried out under general anesthesia via intraperitoneal injection of Ketaminol (45 mg/kg) and Domitor (0.3 mg/kg). Marcain (0.1 mL) was given as a local anesthesia subcutaneously. The rats were placed in a stereotaxic frame and put in a “flat head” position (vertical difference between lambda and bregma < ±0.2 mm). Unilateral lesions of 6-OHDA (3.5 μg/μL, 3μL, 0.3 μL/min) were placed into the MFB to ensure a widespread DA neuron depletion in one hemisphere. The coordinates in SDs were A/P: −4.4, M/L: −1.1, D/V: −7.8 and for nudes, A/P: −3.9, M/L: −1.2, D/V: −7.3. Cells were transplanted 4 weeks after 6-OHDA lesions into the dopamine denervated striatum via two 2 µl deposits in a single tract. For SDs the following coordinates were used: A/P: +0.8, M/L: −2.8, D/V: −4.5/−5.5. For transplantation in athymic rats the following coordinates were used instead: A/P: +1.2, M/L: −2.8, D/V: −4.5/−5.5. For each deposit injection rate of 1μL/min with a 2min diffusion time was used. Rabies virus was injected (3 μL, 0.3 μL/min) one week before the end of the experiment and was placed in the middle of the cell grafting tracts as two deposits of the same D/V coordinates (−4.0/−5.0). Post-surgery, antisedan (0.28 mg/kg) and temgesic (0.04 mg/kg) were given subcutaneously to reverse the anesthesia and serve as an analgesic, respectively.

## Preparation of cells for transplantation

On the day of transplantation, cells were harvested and suspended in Hanks balanced salt solution (HBSS) with 10% DNAse and put on ice until used. 2D reprogrammed cells were dissociated using Accutase (75 µl/cm^2^, 10 min incubation). iDANoids were harvested by gentle pipetting through a cut 1 ml tip. 900.000 hDFs were reprogrammed for transplantation in each animal.

### ΔG-Rabies virus production

EnvA-pseudotyped ΔG-rabies virus was produced according to previously published procedure ^56^. The titer was estimated to 10-30×10^6^ transducing units/mL and was used at a working dilution of 5% for the experiments.

### Immunohistochemistry

At the end of the experiment the rats were administered a lethal dose of anesthesia by intraperitoneal injection of sodium pentobarbital and perfused trans-cardinally with room-temperature 0.9% saline for 3-5min followed by ice-cold 4% PFA (pH 7.4 ± 0.2) for 5 minutes. The brains were manually removed from the skull and were left in 4% PFA at 4 °C overnight. The PFA was removed the next day and replaced with 25% sucrose solution. After 2-3 days, once the brains had sunk in the sucrose solution, they were placed onto a freezing microtome and sectioned at 35 μm thickness in 1:8 series and stored in antifreeze at 4 °C until the start of staining.

All brain sections were stained as free-floating sections in glass vials on a shaker. The sections were washed 3x in 0.1M KPBS, incubated in TRIS-EDTA (pH 9.0) for 30 min at 80 °C to improve antigen binding and washed again 3x. Sections that were DAB-stained were incubated in a quenching solution (10% H_2_O_2_ and 10% methanol in KPBS) for 15 minutes in order to quench endogenous peroxidase activity and washed 3x. Next, the sections were incubated for 1h in blocking solution (5% serum and 0.25% Triton) at RT and were afterwards left to incubate in primary antibodies in blocking solution on a shaker at RT overnight. The following day the sections were washed 2x and put in blocking solution at RT for 30min followed by 1hour in secondary antibodies. For immunofluorescence stainings, the secondary antibodies were fluorophore-conjugated antibodies (1:200 Jackson ImmunoResearch Laboratories) while for DAB stainings biotinylated secondary antibodies were used instead. The immunofluorescent sections were washed 3x, mounted onto chrome alum-gelatin coated slides and coverslipped with PVA-DABCO with added DAPI (1:1000). The DAB sections were washed 3x and incubated in an avidin-biotin complex (ABC Complex) for 1h and washed 3x. The sections were thereafter left to incubate in 0.05% DAB-solution for 2min before adding 20μL of 3% H_2_O_2_ to allow color development. Finally, the sections were washed 3x, mounted onto chrome alum-gelatin coated slides, dehydrated in an ascending order of alcohol, cleared in xylene and coverslipped with DPX mountant.

### Whole brain iDISCO and light sheet microscopy

Two brains were processed for light sheet microscopy using the iDISCO clearing method (Renier et al., 2014) at 8 weeks after cell grafting. The brains were perfused with 2% ice-cold PFA for 5min, followed by 1-hour post-fixation on ice and left stored in PBS. Divided by a midline sagittal cut, each half-brain was processed individually. They were washed in PBS 3x for 30 min, then in 20, 40, 60 and 80% methanol (in PBS), and 100% methanol, for 1 hour each. Next, the brains were incubated in a DCM-methanol mixture (2:1) overnight at RT on a shaker, followed by 2x 30 min washes in 100% methanol and lastly 1 hour at 4 °C. They were bleached in 5% H_2_O_2_ and left on a shaker at 4 °C overnight. Afterwards, they were rehydrated from methanol to 20, 40, 60, 80, 100% PBS, with 30 min for each step and washed in PBS/2% TritonX (PTx.2) 2x for 1 hour at RT, followed by a wash in PBS/0.2% Tween-20 with 10 μg/mL heparin (PTwH) 2x 30 min. Subsequently, the brains were incubated in permeabilization buffer (PTx.2/glycine/DMSO) at 37 °C for 3-5 days on a shaker before being washed 2x 30 min in PTx.2 and incubated with primary antibody in PTwH, 5% DMSO/3% Serum (NDS) at 37 °C for 10 days: 1:500 goat polyclonal anti-mCherry (SciGen, Ab0040). Thereafter, the brains were washed in PTwH for 10, 15, 20 min, then every hour during the day, then incubated at 37 °C with donkey Cy5 secondary antibody 1:500 in PTwH/3% normal donkey serum (NDS) for the next 10 days. Following this, they were washed in PTwH for 10, 15, 20 min, then every hour during the day, and left overnight. Then they were dehydrated to 20, 40, 60, 80, then 100% methanol, 1 hour for each step, followed by 100% methanol overnight. The next day they were put in DCM-methanol (6:4) on a shaker overnight and the day after they were incubated in 100% DCM 15 min twice, then changed to DiBenzyl Ether (DBE) for clearing. All solutions were filtered before use and 0.02% NaN_3_ was added to all stock solutions to prevent microbial growth.

The cleared hemispheres were then imaged on an Ultra Microscope II (LaVision Biotec) equipped with an sCMOS camera (Andor Neo, model 5.5-CL3) and 4x or 12x objective lenses (LaVision LVMI-Fluor 4x/0.3 or 12x/0.53 MI Plan). Two laser configurations with following emission filters: 525/50 for endogenous background and AlexaFluor 488, and 680/30 for AlexaFluor 647 were used. Image stacks were acquired with ImspectorPro64 (LaVision Biotec) utilizing 5 μm z-steps. All samples were imaged in a chamber filled with DBE. Several stacks (mosaic acquisition) were taken with 10% overlap to cover each hemisphere and the stacks were stitched to visualize the brain in 3D with Arivis Vision 4D 3.01 (Arivis AG). The movies were then compiled in Final Cut Pro 10.4.3 (Apple Inc.).

### Whole cell patch clamp recordings in acute brain slices

Animals were decapitated under anesthesia with Isoflorane (Baxter, Germany). The brain was quickly removed from the skull and placed in an ice-cold oxygenated (carbogen, 5% CO2, 95% O2) N-methyl-d-glucamine (NMDG)-HEPES artificial cerebrospinal fluid (aCSF) containing the following (in mM): 92 NMDG, 2.5 KCl, 1.25 NaH2PO4, 30 NaHCO3, 20 HEPES, 25 glucose, 2 thiourea, 5 Na-ascorbate, 3 Na-pyruvate, 0.5 CaCl2·2H2O, and 10 MgSO4·7H2O. We titrated the pH to 7.3–7.4 with 7 mL ± 0.2 mL of 37% hydrochloric acid, osmolarity ranging from 300 to 305 mOsm/kg. Acute brain slices containing the dorsolateral striatum were prepared at 275μm thickness using a vibratome (Leica VT1200 S, Wetzlar, Germany). Slices were incubated for 25 min at 35 °C and then transferred at room temperature into a holding chamber. The recordings were carried out with continuous perfusion of aCSF (1 ml/min) containing (in mM): 119 NaCl, 2.5 KCl, 1.3 MgSO4, 2.5 CaCl2, 25 Glucose and 26 NaHCO3, and gassed with 95% O2–5% CO2 at room temperature (pH ∼7.4, 305 mOsm/kg). The cells were visualized using a fixed-stage Olympus Microscope (BX51WI) coupled with an IR-CCD camera and a 40X water immersion objective. Patch pipettes (4–6 MOhm resistance) were pulled from standard borosilicate glass (BF150-86-10, O.D.: 1.5mm, I.D.: 0.86 mm, with filament, Sutter Instrument, Novato, CA, USA) by using a micropipette puller (p1000, Sutter Instrument, Novato, CA, USA). Recording pipettes were filled with an intracellular solution containing (in mM): 130 (K)Gluconate, 10 KCl, 0.2 EGTA, 10 HEPES, 4 (Mg)ATP, 0.5 (Na)GTP, 10 (Na)Phosphocreatine (pH 7.25, 296 mOsm). Whole-cell patch-clamp recordings were obtained using a Multiclamp 700B amplifier and pClamp 10.4 data acquisition software (Axon Instruments, Molecular Devices, USA). The access resistance was monitored throughout the recording, and cells were discarded if the access resistance was over 35 MOhm. Resting membrane potential (RMP) was measured right after breaking into the cell. The capacitance of the cell was directly obtained from pClamp 10.4 data acquisition software. The input resistance (Ri) was calculated in the current clamp from a 20pA hyperpolarizing current injection for 500 ms. Action potential threshold, peak amplitude after hyperpolarization (AHP) and halfwidth were measured from the first observed spike evoked by the rheobase current injection step. Voltage responses to current injection from −20 to +35 pA with a delta of 5 pA for 500 ms were recorded at a holding potential of −70 mV. Activation of voltage-dependent sodium and potassium channels was carried out using increasing depolarizing voltage steps from −70 mV to + 40 mV with a delta of 10 mV for 100ms. Sodium channels were selectively blocked using tetrodotoxin (TTX 1µM, Tocris, Bristol, United Kingdom) by adding to the extracellular solution in some recordings. Data were analyzed offline using Clampfit 10.3 (Axon Instruments, Molecular Devices, USA) and Igor Pro 8.04 (Wavemetrics, Portland, OA, USA) combined with the NeuroMatic package.

## Acknowledgments

This work was supported by funding from the New York Stem Cell Foundation, European Research Council (ERC) under ERC Grant Agreement 771427, European Union–funded project NSC-Reconstruct (European Union, H2020, GA no 874758, 2020-23), Swedish Research Council (2021-00661, 2021-02967, and 2021-01839), Swedish Parkinson Foundation (Parkinsonfonden), Swedish Brain Foundation, Konung Gustaf V:s och Drottning Victorias Frimurarestiftelse, Olle Engkvist Foundation (213-0229), Anna-Lisa Rosenberg Foundation, Royal Physiographic Society in Lund (43202), and Strategic Research Area at Lund University Multipark. Mette Habekost was supported by the Lundbeck Foundation Postdoc Fellowship (R347-2020-2522).

The authors would like to thank Michael Sparrenius for the help with animal work; Ulla Jarl and Bengt Mattsson for providing help with sample preparation, imaging, and image reconstruction for iDISCO of the whole rat brains; Sol da Rocha Baez for the help with virus production; Natalia Avaliani from Electrophysiology core facility at Lund Stem Cell Center for the help with *ex vivo* electrophysiological measurements. Lund University Bioimaging Centre (LBIC) is gratefully acknowledged for providing experimental resources for confocal microscopy. Methylation profiling was performed by the SNP&SEQ Technology Platform in Uppsala (www.genotyping.se). The facility is part of the National Genomics Infrastructure supported by the Swedish Research Council for Infrastructures and Science for Life Laboratory, Sweden.

## Contributions

J.K., F.N., M.B., A.F., M.H., and M.P. designed the study. J.K., F.N., K.L., and M.B. carried out experiments and analysis. A.B., and E.C.P. carried out the electrophysiology experiments and analysis. R.A.B., D.R.O, and A.H. contributed new reagents/analytic tools. P.S. and M.H. performed the bioinformatics analysis. J.K. and M.P. wrote the first draft of the paper.

## Notes

### Competing Interest Statement

The authors have declared no competing interest.

