## Supplemental information for "Injectable 3D microcultures enable intracerebral transplantation of mature neurons directly reprogrammed from patient fibroblasts"

Supplementary Figures:

Supplementary Figure 1
Supplementary Figure 2
Supplementary Figure 3
Supplementary Figure 4
Supplementary Figure 5

Supplementary Tables:

Supplementary Table 1
Supplementary Table 2

Supplementary Movies:

Supplementary Movie 1
Supplementary Movie 2

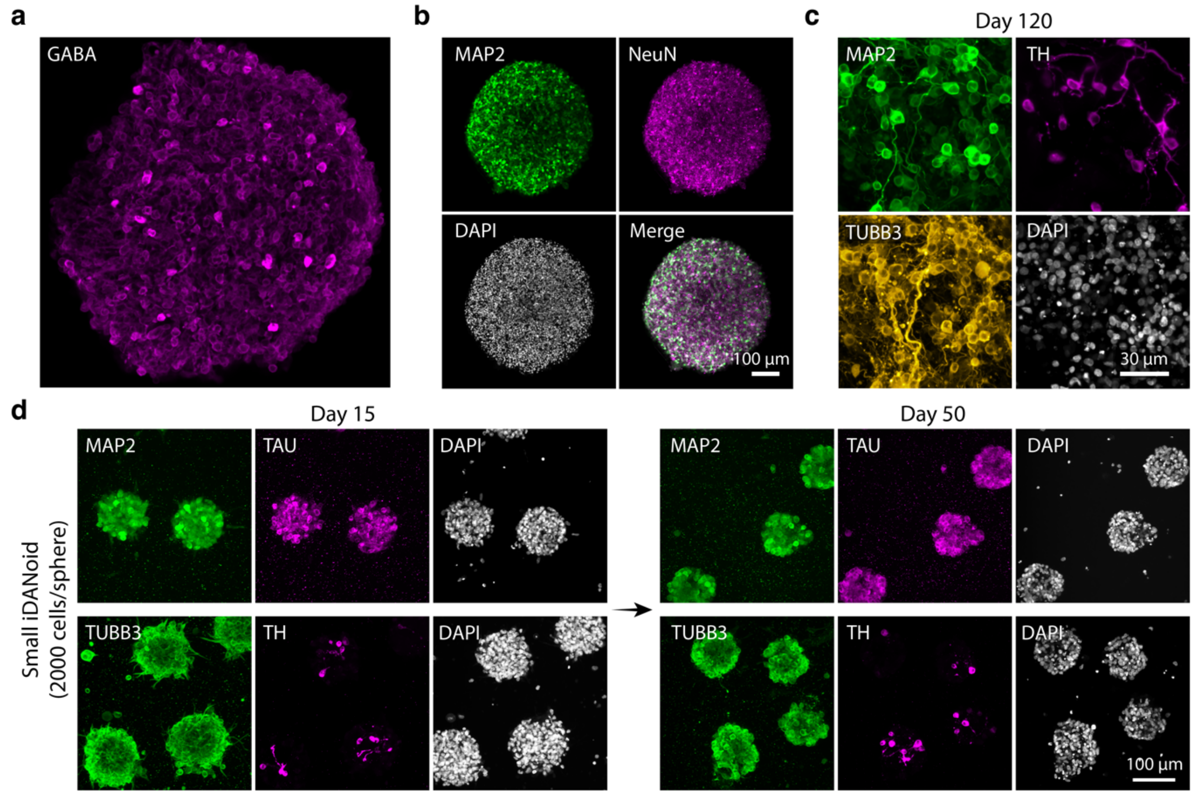


**Supplementary figure 1**

Additional immunocytochemical analysis of iDANoids. **a**, Fluorescence confocal image showing the presence of GABAergic neurons inside iDANoids at day 30. **b**, Overview fluorescence image showing the presence of mature neuronal marker NeuN in iDANoids at day 50. **c**, Fluorescence image showing directly reprogrammed neurons inside iDANoids after 120 days in culture. **d,** Fluorescent confocal images of neuronal (MAP2, TAU, TUBB3) and DA (TH) markers in iDANoids at day 15 and day 50.

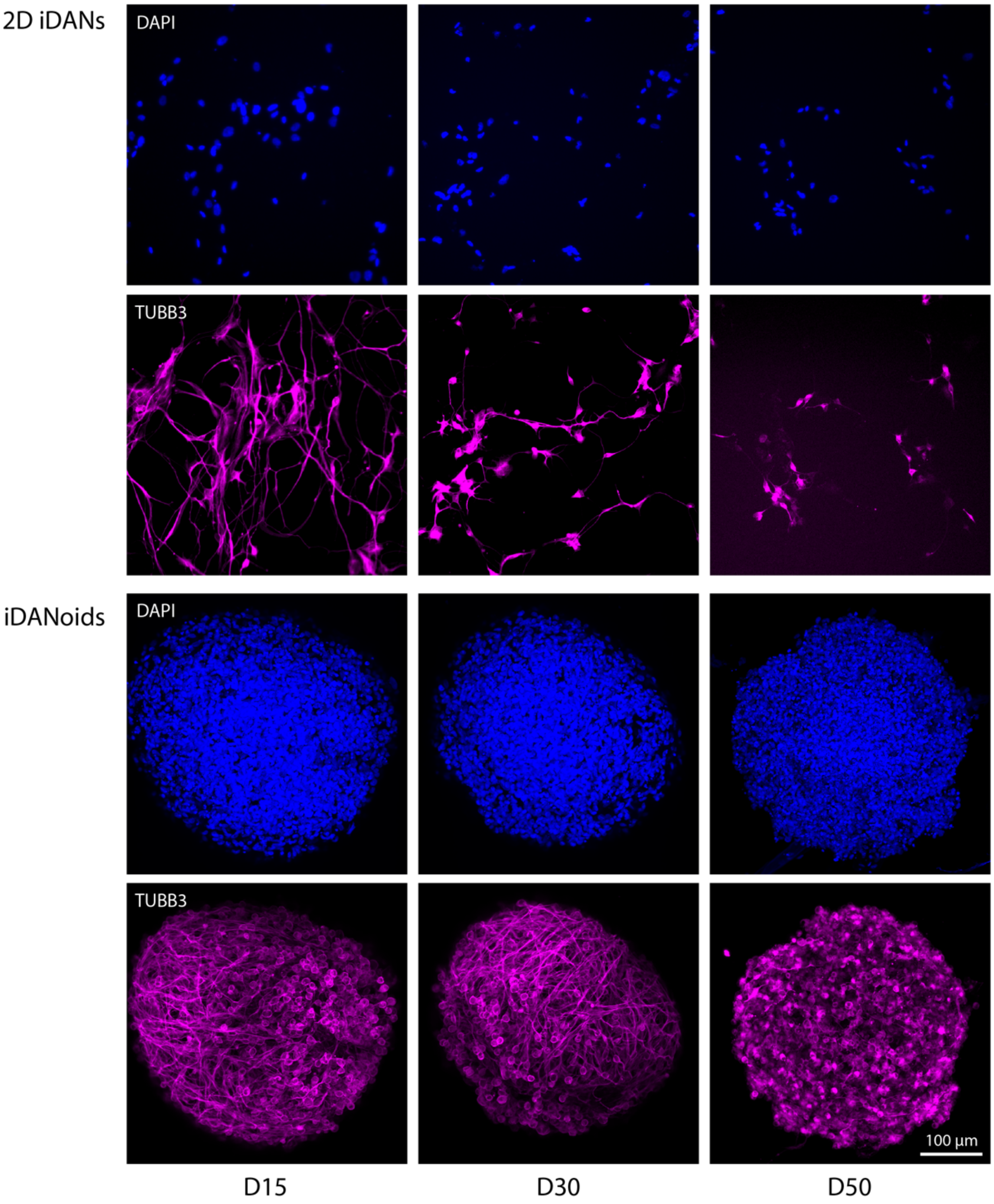


**Supplementary figure 2**

Representative confocal images showing nuclei (DAPI) and neurons (TUBB3) for three different timepoints and for reprogramming in 2D and 3D (iDANoids).

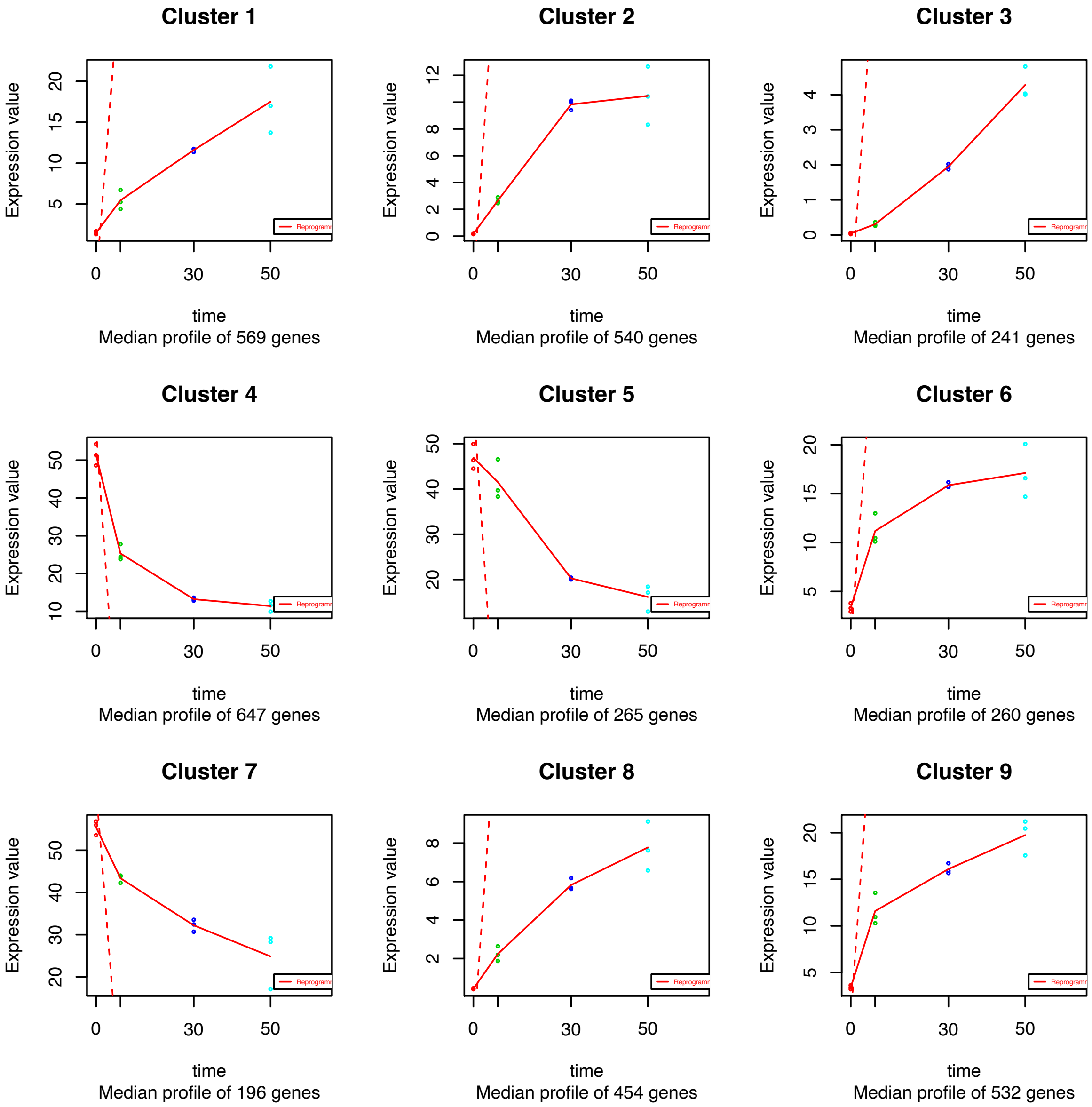


**Supplementary figure 3**

Results of regression-based maSigPro analysis showing 9 clusters of genes with significant expression profile difference along the experimental time course (D0, D7, D30, D50).

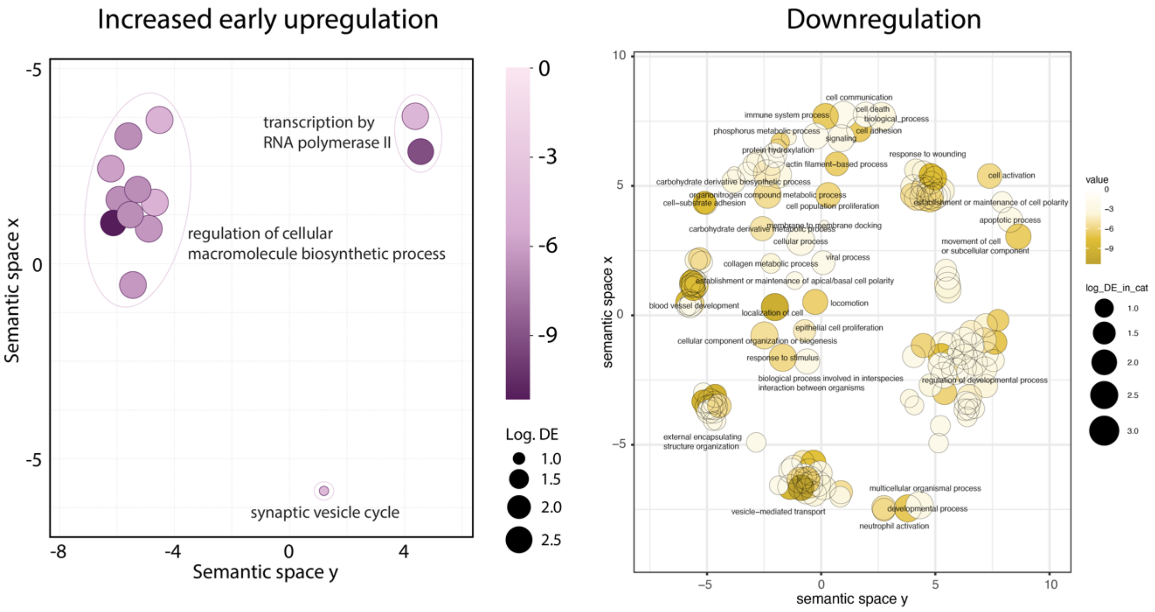


**Supplementary figure 4**
Gene ontology (GO) analysis of biological processes overrepresented in gene sets with early upregulation (clusters 6 and 9 from maSigPro analysis in Figure 3d) and in downregulated gene sets (clusters 4, 5, and 7 from maSigPro analysis in Figure 3d).

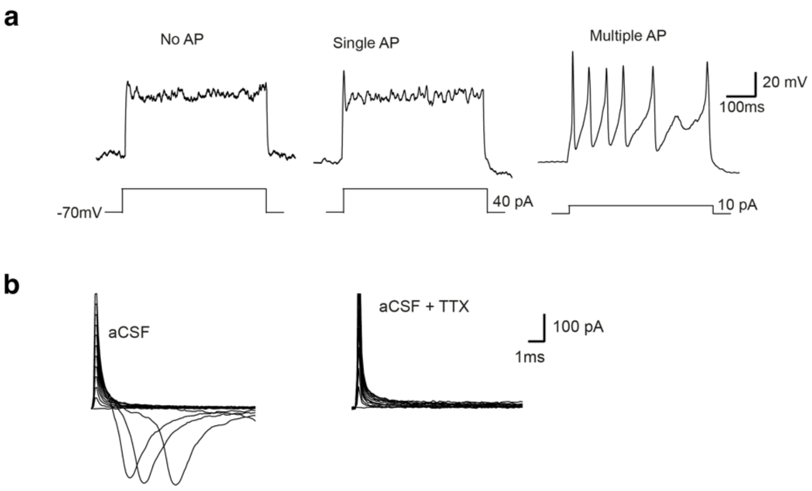


**Supplementary figure 5**

Complementary data to results from whole-cell patch-clamp recordings on coronal acute brain slices presented in Figure 7. **a,** Representative traces of voltage responses evoked by +40 pA hyperpolarizing current for cells firing none, single and multiple action potentials (APs). **b,** Inward sodium current recorded in the presence of tetrodotoxin (1µM, TTX).

Supplementary Table 1 - **: List of primers used for RT-qPCR analysis in this study.**

| **Primers** | | **Sequence (5’-3’)** | **Full gene name** |
|---|---|---|---|
| *AADC* | F | GGGGACCACAACATGCTGCTCC | DOPA decarboxylase (DDC) |
| | R | AATGCACTGCCTGCGTAGGCTG | |
| *ACTB* | F | CCTTGCACATGCCGGAG | Beta-actin |
| | R | GCACAGAGCCTCGCCTT | |
| *DAT* | F | CACTGCAACAACTCCTGGAA | Dopamine transporter (SLC6A4) |
| | R | AAGTACTCGGCAGCAGGTGT | |
| *GAPDH* | F | TTGAGGTCAATGAAGGGGTC | Glyceraldehyde-3-phosphate dehydrogenase |
| | R | GAAGGTGAAGGTCGGAGTCA | |
| *PITX3* | F | GGAGGTGTACCCCGGCTACTCG | Paired-like homeodomain 3 |
| | R | GAAGCCAGAGGCCCCACGTTGA | |
| *SYN* | F | CCCGTGGTTGTGAAGATGGGGC | Synapsin 1 |
| | R | TGCCACGACACTTGCGATGTCC | |
| *TH* | F | CGGGCTTCTCGGACCAGGTGTA | Tyrosine hydroxylase |
| | R | CTCCTCGGCGGTGTACTCCACA | |

Table 1: List of primers used for RT-qPCR analysis of reprogrammed iDANoids.

Supplementary Table 2 – List of primary antibodies used in this study.

| Marker | Specificity | Source (cat. no.) | Dilution |
|---|---|---|---|
| GABA | Rabbit | Sigma (A2052) | 1:2000 |
| GFP | Chicken | Abcam (AB13970) | 1:1500 |
| hNCAM | Mouse | Santa Cruz (sc 106) | 1:500 |
| HuNu | Mouse | Millipore (MAB 1281) | 1:200 |
| MAP2 | Chicken | Abcam (ab5392) | 1:5000 |
| mCherry | Goat | Sicgen (AB0040-200) | 1:500 |
| NeuN | Rabbit | Merck Millipore (ABN78) | 1:1000 |
| TAU | Rabbit | DAKO (A0024) | 1:1000 |
| TH | Rabbit | Merck Millipore (AB152) | 1:1000 |
| TUBB3 | Mouse | BioLegend (801202) | 1:500 |
| Vimentin | Mouse | DAKO (M0725) | 1:50 |

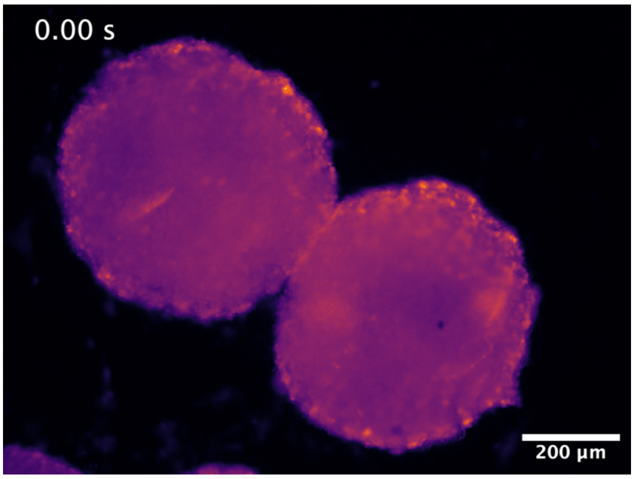


**Supplementary movie 1** Timelapse of confocal fluorescence images using Calbryte AM dye to show a $Ca^{+}$ response of D30 iDANoids to KCl stimulation.
